# Histone H1 Promotes Silencing of Unintegrated HIV-1 DNA

**DOI:** 10.1101/2025.11.07.687224

**Authors:** Jiaqi Zhu, Gary Z. Wang, Hugo D. Pinto, Sean E. Healton, Laxmi N. Mishra, Arthur I. Skoultchi, Stephen P. Goff

## Abstract

In eukaryotic cells, genomic DNA is packaged into chromatin with nucleosomes formed by core histones H2A, H2B, H3, and H4, and further stabilized by the linker histone H1. During the early stages of retroviral infection, such as with murine leukemia virus (MLV) and human immunodeficiency virus type 1 (HIV-1), host core and H1 histones are rapidly deposited onto unintegrated viral DNAs upon nuclear entry. These unintegrated viral DNAs are transcriptionally silenced through histone post-translational modifications (PTMs), including high levels of H3K9 trimethylation and low levels of H3 acetylation. Linker histone H1 is closely associated with chromatin compaction and histone PTMs, suggesting a potential role in regulating retroviral DNA fate. In this study, we demonstrate that simultaneous knockdown of four somatic H1 variants (H1.2, H1.3, H1.4, and H1.5) in K562 cells reverses the silencing of unintegrated HIV-1 DNA, resulting in increased viral expression. Notably, this effect was specific to HIV-1, as the same H1 depletion did not alter the silencing of MLV unintegrated DNA. These results reveal distinct roles of H1 in regulating HIV-1 and MLV unintegrated DNA expression.

## Introduction

Retroviral replication begins with the conversion of the single-stranded RNA genome into double-stranded DNA, catalyzed by the viral reverse transcriptase shortly after infection (1). The newly synthesized linear viral DNA is flanked by long terminal repeats (LTRs) and is assembled into a nucleoprotein structure known as the preintegration complex (PIC) (2, 3). The PIC protects the viral DNA and mediates its nuclear import, though the pathways differ across retroviral genera. For example, the PICs of gammaretroviruses such as Moloney murine leukemia virus (MLV) can only access host chromatin during mitosis, when the nuclear envelope breaks down (4, 5). In contrast, lentiviruses such as HIV-1 are able to infect non-dividing cells, as their PICs are actively transported through the nuclear pore complex via interactions between capsid proteins (CA) and specific nucleoporins (6–9). Once inside the nucleus, viral DNA can follow two fates. It may integrate into the host genome through the action of viral integrase (IN), forming the provirus that serves as a template for both viral mRNA transcription and genomic RNA production. Alternatively, unintegrated viral DNAs can persist as episomes. Some are circularized through homologous recombination, generating one-LTR circles (10), while others are processed by non-homologous end joining (NHEJ) to form two-LTR circles (2-LTR) (11). The integrated proviruses exhibit robust viral gene expression, while integration-deficient mutants show poor expression and gradually decay (12, 13). Both core histones and linker histone H1 are rapidly deposited onto unintegrated viral DNAs after nuclear entry, which display high levels of the repressive mark H3K9 trimethylation and low levels of H3 acetylation, resulting in transcriptional silencing (14, 15). The magnitude of the silencing of unintegrated DNA relative to integrated DNA varies widely among cell lines, and is particularly dramatic in lymphoid lines such as the K562 line (16). Importantly, the transcriptional silence of unintegrated viral DNA can be partially relieved by treatment with histone deacetylase (HDAC) inhibitors, highlighting the role of histone modification in restricting viral gene expression (17). Furthermore, genome-wide CRISPR knockout screens identified several key host factors involved in silencing unintegrated MLV DNA, including the DNA-binding protein NP220, all three subunits of the HUSH complex, the H3K9 methyltransferase SETDB1, and selected HDACs (18). Similarly, the histone chaperones CHAF1A and CHAF1B, as well as the SMC5/6 complex, have been implicated in the silencing of unintegrated HIV-1 DNAs (19–21). Notably, the various host factors are not all involved in the silencing of both MLV and HIV-1, but often only impact one or the other virus. These findings strongly suggest that histones, particularly histone modifications, play an essential role in the silencing of unintegrated retroviral DNA.

In eukaryotic cells, chromatin is organized into nucleosomes, each consisting of ~147 bp of DNA wrapped around an octamer of core histones H2A, H2B, H3, and H4 (22, 23). H1 linker histones bind to DNA at the entry and exit sites of nucleosomes, stabilizing nucleosome positioning and promoting higher-order chromatin folding (24). Higher eukaryotes encode a diverse repertoire of H1 linker histone variants, with 11 distinct H1 genes identified in both mice and humans. Among them, H1.1–H1.5 are somatic subtypes that are broadly expressed in a replication-dependent manner (25, 26). In contrast, H1.0 and H1x are expressed independently of the cell cycle (27, 28). Additional specialized subtypes are found in germ cells, including H1oo in oocytes and H1t, H1T2, and H1LS1 in spermatids or spermatocytes (29–31). The fundamental role of H1 is to work together with core histones to stabilize nucleosome architecture and higher-order chromatin folding, and to influence the nucleosome repeat length (NRL) (32). Beyond this structural role, H1 variants can regulate gene expression by modulating transcription factor binding and shaping the epigenetic landscape through effects on both histone and DNA modifications (24, 26, 33, 34).

Despite the central role of histone-based repression in controlling unintegrated viral DNA, the specific contributions of H1 remain poorly defined. In this study, we investigate the role of H1 in the regulation of unintegrated retroviral DNA, focusing on HIV-1 and MLV, with the aim of defining how H1 contributes to the viral DNA silencing.

## Results

### H1 knockdown affects expression of HIV unintegrated DNAs

To investigate whether histone H1 influences expression from unintegrated HIV-1 DNAs, the expression of four H1 subtypes (H1.2, H1.3, H1.4, and H1.5) was transiently knocked down in K562 cells using the CRISPRi system. Control cells and H1 knockdown (H1-KD) cells were then infected with pNL4-3.GFP.R^−^.E^−^ HIV reporter viruses carrying either wild-type integrase (IN-WT) or the integration-deficient mutant (IN-D64A). In control cells, IN-WT infection resulted in high GFP expression, while IN-D64A infection resulted in dramatically reduced GFP expression. Treatment with the histone deacetylase inhibitor TSA restored expression to high levels. In contrast, in H1-depleted cells, infection with IN-D64A resulted in high level GFP expression, comparable to infection with IN-WT virus, or with TSA-treated cells (Fig. 1A). H1 depletion had no significant impact on expression from the IN-WT virus. The same analysis was performed using MLV reporter viruses expressing a GFP gene delivered in either IN-WT or an integration-deficient mutant (IN-D184A) virions. As expected, expression delivered by the integration-deficient mutant was heavily silenced, but unexpectedly, H1 depletion did not restore expression from the IN-D184A mutant. Instead, the only effect was a modest reduction in IN-WT expression observed in H1-depleted cells (Fig. 1B). Analysis of viral DNAs by PCR revealed no significant difference in total viral DNA or 2-LTR circle levels between control and H1-KD cells infected with either IN-WT or IN-D64A HIV (Fig. 1C). Similarly, in MLV-infected cells, total viral DNA levels were comparable between the two cell types, although 2-LTR circles derived from the IN-D184A mutant were slightly reduced in H1-KD cells (Fig. 1C). Collectively, these findings suggest that histone H1 plays distinct roles in regulating expression from unintegrated HIV and MLV DNAs – a major role in silencing unintegrated HIV-1 DNA and no detectable effect on MLV DNA.

**Figure 1.**
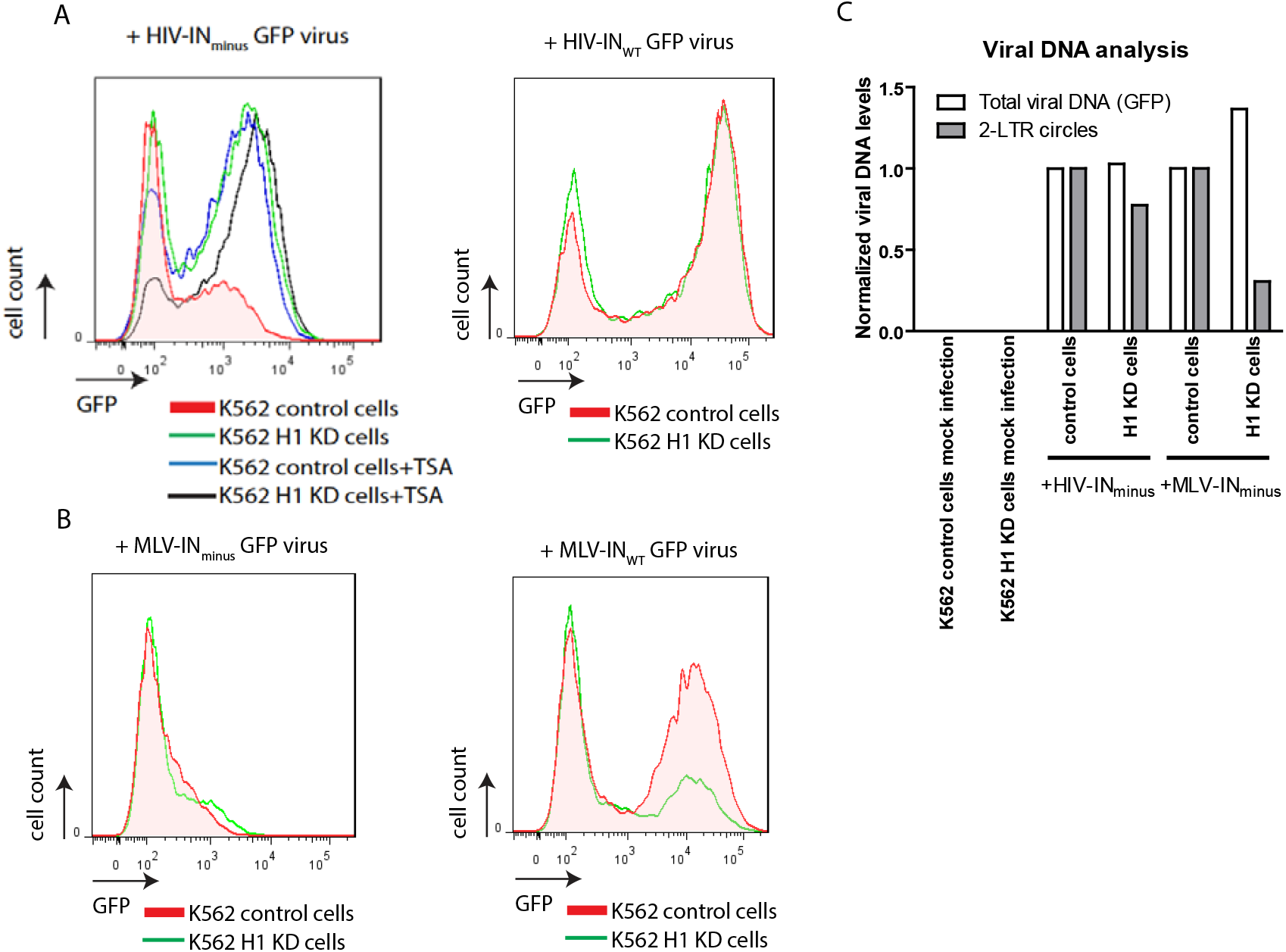
H1 knockdown affects expression of HIV unintegrated DNAs. A) Control and H1-KD K562 cells were infected with equal volumes of pNL4-3.GFP.R^−^.E^−^ reporter viruses carrying either wild-type integrase (IN-WT) or the integration-deficient mutant (IN-D64A). Infected cells were cultured in the presence or absence of TSA. At 48 h post-infection, GFP fluorescence was analyzed by flow cytometry, and representative histograms showing showing the distribution of GFP fluorescence are presented. B) Control and H1-KD K562 cells were infected with equal volumes of pNCA-GFP reporter viruses carrying either wild-type integrase (IN-WT) or the integration-deficient mutant (IN-D184A). At 48 h post-infection, cells were analyzed by flow cytometry, and representative histograms showing the distribution of GFP fluorescence are presented. C) qPCR analysis of total viral DNA (GFP) and unintegrated DNA (2-LTR circles) from cells collected in A) and B).

### H1 knockdown reduces expression of H1 variants in iK562 cells

To extend these observations, utilizing alternative methods of H1 depletion that avoid acute transfection, we generated inducible silencing constructs stably integrated into the genome. Doxycycline (Dox)-inducible H1 knockdown K562 cells were generated by introducing doxycycline-inducible dCas9-KRAB together with sgRNAs targeting all four H1 variants (H1.2, H1.3, H1.4, and H1.5), or a scrambled control sgRNA (see Methods). Scrambled control (iK562-Scr) and histone H1 knockdown (iK562-H1-KD) cells were treated with 3 μg/ml Dox for 7 days. Cells were collected at days 0, 3, 5, and 7, and histones were extracted for western blotting. Histone H3, which remains relatively constant across conditions, was used as a loading control. At day 0, the expression of H1.2, H1.3, H1.4, H1.5, and H1.x was comparable between iK562-Scr and iK562-H1-KD cells. Upon Dox induction, H1.2 decreased to around 40% of control levels, while H1.3, H1.4, and H1.5 were reduced by approximately 50–60% by day 5 and remained low through day 7. In contrast, H1.x displayed a progressive increase following induction, suggesting a compensatory mechanism in response to the loss of other H1 variants (Fig. 2A, B). To assess the H1 content within total histones, samples isolated on day 5 were analyzed by SDS–PAGE followed by Coomassie staining. A substantial reduction in total H1 variants was detected in iK562-H1-KD cells, while the levels of the four core histones (H2A, H2B, H3, and H4) remained comparable between knockdown and control samples (Fig. 2C). To quantify the proportion of H1 within total histones, RP-HPLC was conducted and the results showed that the overall abundance of H1 histones in knockdown cells was reduced to about 25% of the levels observed in control cells (Fig. 2D, E). Together, these findings demonstrate that doxycycline induction effectively reduces multiple H1 variants in iK562-H1-KD cells, while the core histones remain unaffected. The observed upregulation of H1.x likely represents a compensatory response to maintain chromatin organization in the context of H1 depletion.

**Figure 2.**
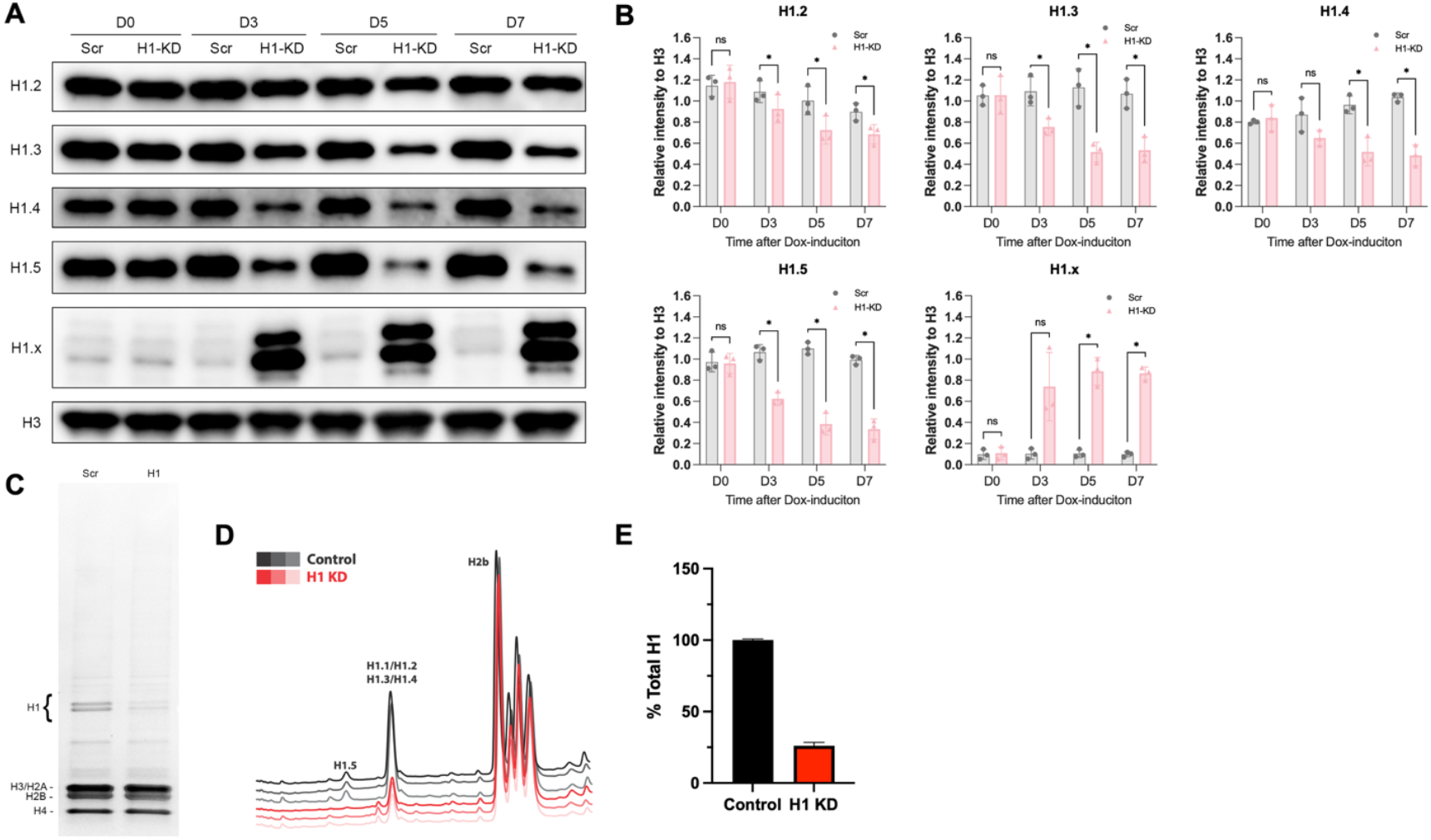
Histone H1 knockdown reduces expression of H1 variants in iK562 cells. A) iK562-Scr and iK562-H1-KD cells were treated with Dox and collected at days 0, 3, 5, and 7. Histones extracted from cells were analyzed by Western blotting. Representative blots are shown, probed with antibodies against individual H1 variants. Histone H3 was used as a loading control. B) Quantification of H1 variant protein levels from the Western blots in (A), shown as the ratio of H1 variant band intensity to H3. Error bars indicate mean ± SD; n = 2 independent experiments. C) Coomassie-stained gel showing histones isolated from iK562-Scr and iK562-H1-KD cells at day 5 after Dox induction. D) RP-HPLC analysis of histones from iK562-Scr and iK562-H1-KD cells at day 5 after Dox induction. n = 3 independent experiments. E) Quantification of H1 as a percentage of total histones from iK562-Scr and iK562-H1-KD cells at day 5 after Dox induction. Error bars indicate mean ± SD; n = 3 independent experiments.

### H1 knockdown restores expression of integration-deficient HIV-1 in different reporter constructs

Based on our previous observation that H1 knockdown reaches its maximum effect by day 5 after induction, iK562 cells were infected at this time point with equal volumes of pNL4-3.mCherry.R^−^.E^−^ reporter viruses, which express mCherry in place of *nef* from a spliced RNA, or treated with medium alone as a mock control. The viruses carried either wild-type integrase (IN-WT) or the integration-deficient mutant (IN-D64A). At 48 h post-infection, the percentage of mCherry-positive cells was quantified by flow cytometry. In control iK562-Scr cells, IN-D64A infection resulted in an approximately 11.8-fold reduction in mCherry-positive cells compared with IN-WT. In contrast, in iK562-H1-KD cells, IN-D64A infection showed only a 1.9-fold reduction relative to IN-WT (Fig. 3A, B), demonstrating that H1 depletion restores reporter expression from unintegrated viral DNAs. No mCherry-positive cells were detected in the mock group. The mean fluorescence intensity (MFI) was markedly lower in cells infected with IN-D64A than in those infected with IN-WT. MFI values were comparable between iK562-Scr and iK562-H1-KD cells, indicating that H1 depletion increases the number of positive cells without enhancing the per-cell reporter expression level (Fig. 3C). To assess whether this difference reflected altered viral DNA levels, total DNA was extracted at 48 h post-infection. Total viral DNA from IN-D64A infections was slightly lower than that from IN-WT infections, but this decrease occurred similarly in both iK562-Scr and iK562-H1-KD cells (Fig. 3D). In contrast, 2-LTR circle DNAs were significantly increased in IN-D64A infections in both cell types, consistent with the accumulation of unintegrated viral DNA (Fig. 3D). No viral DNA or 2-LTR circles were detected in mock samples. Comparable results were obtained using pLVX-mCherry reporter viruses, which express mCherry from the CMV promoter via an unspliced RNA, carrying IN-WT or IN-D64A. Both HIV-1 reporter viruses were constructed without Vpr expression to avoid induction of cell cycle arrest (35). In iK562-Scr cells, mCherry expression from IN-D64A virus was strongly suppressed, whereas in iK562-H1-KD cells expression was restored to levels approaching those of IN-WT (Fig. 3E, F). Interestingly, the MFI did not differ substantially between IN-D64A– and IN-WT–infected cells, and both MFI values and viral DNA levels followed similar patterns in the two cell lines (Fig. 3G, H). Collectively, these findings demonstrate that H1 knockdown restores expression from unintegrated HIV-1 DNA without altering viral DNA abundance. Moreover, because pNL4-3.mCherry.R^−^.E^−^ and pLVX-mCherry express mCherry from spliced and unspliced reporter transcripts respectively, the rescue of expression does not depend on viral RNA splicing.

**Figure 3.**
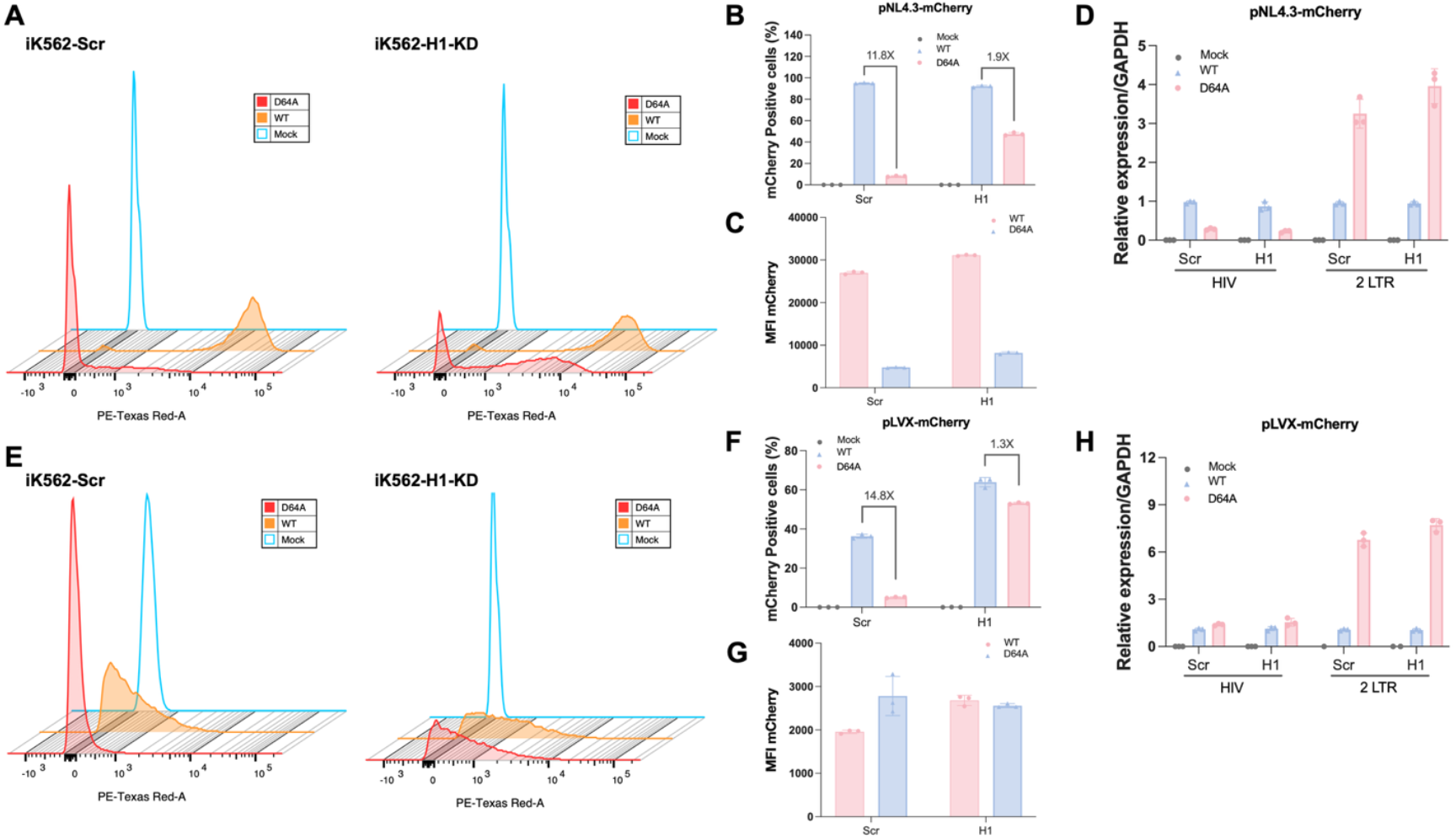
H1 knockdown restores expression from integration-deficient HIV in iK562 cells. A) iK562-Scr and iK562-H1-KD cells were treated with Dox and infected with equal volumes of pNL4-3.mCherry.R^−^.E^−^ carrying either wild-type integrase (IN-WT) or the integration-deficient mutant (IN-D64A), or treated with medium alone as a mock control. At 48 h post-infection, cells were analyzed by flow cytometry, and representative histograms showing the distribution of mCherry fluorescence are presented. B) Percentage of mCherry-positive cells from the flow cytometry in (A). Numbers above the bars indicate the fold difference in mCherry-positive cells between iK562-Scr and iK562-H1-KD. Error bars indicate mean ± SD; n = 3 independent experiments. C) The MFI of mCherry-positive cells from the flow cytometry in (A). Error bars indicate mean ± SD; n = 3 independent experiments. D) qPCR analysis of total viral DNA (HIV) and unintegrated DNA (2-LTR circles) from cells collected in (A). Error bars indicate mean ± SD; n = 3 independent experiments. E) iK562-Scr and iK562-H1-KD cells were treated with Dox and infected with pLVX-mCherry viruses carrying either IN-WT or IN-D64A, or treated with medium alone as a mock control. Cells were analyzed by flow cytometry at 48 h post-infection, and F) representative histograms of mCherry fluorescence are shown. G) Quantification of mCherry-positive cells from the flow cytometry in (D). Numbers above the bars indicate the fold difference in iK562-Scr relative to iK562-H1-KD. Error bars indicate mean ± SD; n = 3 independent experiments. H) The MFI of mCherry-positive cells from the flow cytometry in (E). Error bars indicate mean ± SD; n = 3 independent experiments. I) qPCR analysis of total viral DNA and 2-LTR circles from cells collected in (D). Error bars indicate mean ± SD; n = 3 independent experiments.

### H1 knockdown fails to relieve silencing of integration-deficient MLV in multiple reporter constructs

To determine whether the effect of H1 depletion on unintegrated viral expression extends beyond HIV-1, iK562 cells were infected on day 5 after doxycycline induction with MLV-mCherry reporter viruses, in which mCherry is produced from a spliced transcript replacing the env gene, carrying either wild-type integrase (IN-WT) or the integration-deficient mutant (IN-D184A), with treatment of medium alone as a mock control. At 48 h post-infection, IN-D184A infection resulted in an approximately 41.5-fold reduction in the number of mCherry-positive cells compared to IN-WT in iK562-Scr controls. Although this silencing was less pronounced in H1-depleted cells (~6.1-fold), expression remained strongly suppressed compared to IN-WT (Fig. 4A, B). The MFI was substantially reduced in IN-D184A–infected cells relative to IN-WT infection in both cell lines (Fig 4 C). Analysis of viral DNA revealed no major difference in total DNA levels between Scr and H1-KD cells (Fig. 4D). As expected, 2-LTR circles accumulated in IN-D184A infections, but this increase did not correlate with restored expression (Fig. 4D). Similar results were obtained using a second MLV reporter construct, pNCA-mCherry, in which mCherry is expressed from a spliced RNA substituting for gag-pol. Expression from the IN-D184A mutant remained largely silenced in H1-depleted cells, showing only a minor increase relative to controls (Fig. 4E, F). MFI values were consistent between the two cell lines (Fig. 4G). Although total viral DNA and 2-LTR circles were slightly elevated in H1-KD cells, these changes were not accompanied by meaningful reactivation of reporter expression (Fig. 4H). These findings indicate that, in contrast to HIV-1, H1 knockdown is insufficient to relieve transcriptional silencing of unintegrated MLV DNAs.

**Figure 4.**
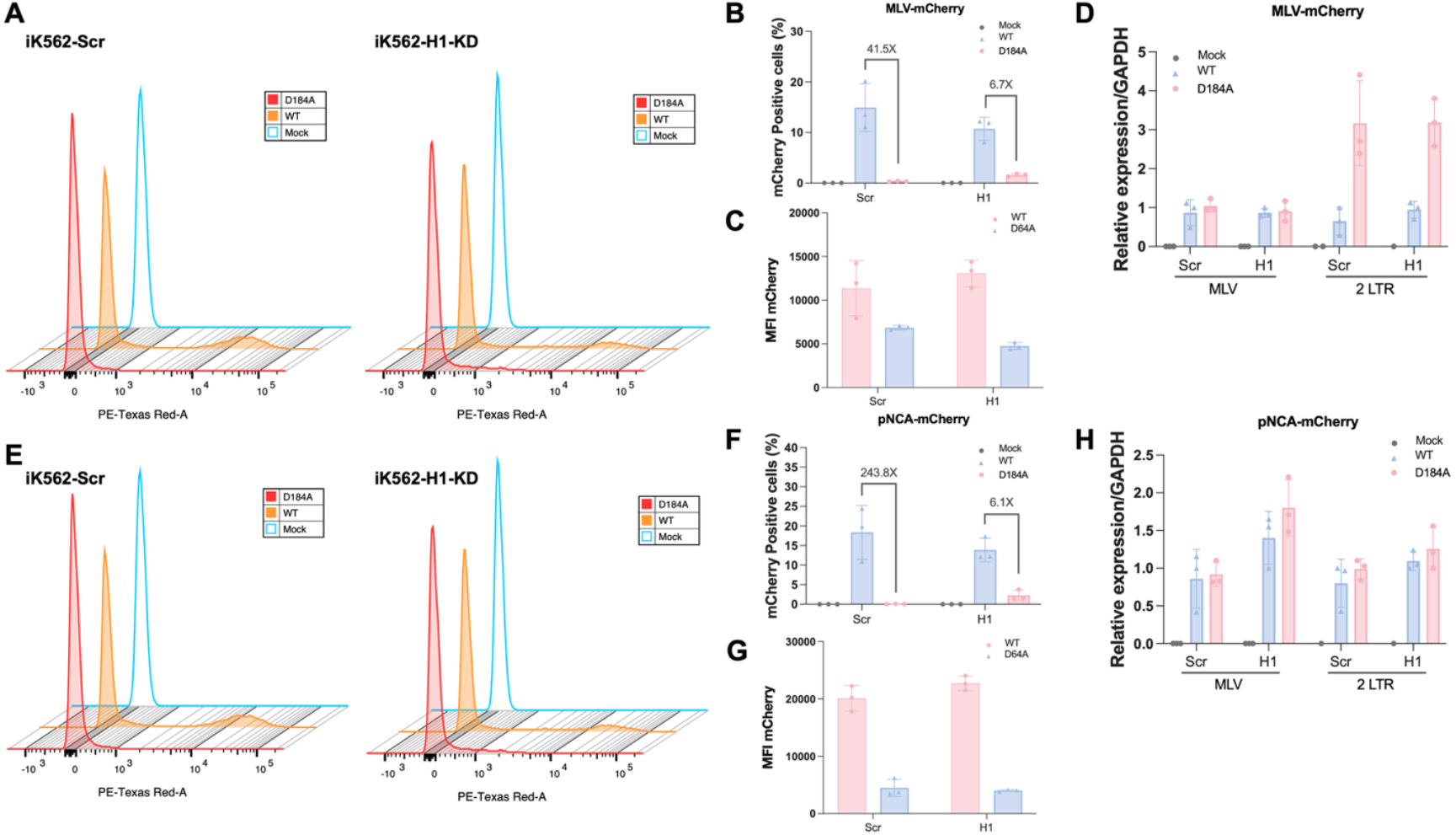
Histone H1 Knockdown Fails to Relieve Silencing of Integration-Deficient MLV in iK562 Cells. A) iK562-Scr and iK562-H1-KD cells were treated with Dox and infected with equal volumes of MLV-mCherry carrying either wild-type integrase (IN-WT) or the integration-deficient mutant (IN-D184A), or treated with medium alone as a mock control. At 48 h post-infection, cells were analyzed by flow cytometry, and representative histograms showing the distribution of mCherry fluorescence are presented. B) Percentage of mCherry-positive cells from the flow cytometry in (A). Numbers above the bars indicate the fold difference in mCherry-positive cells between iK562-Scr and iK562-H1-KD. Error bars indicate mean ± SD; n = 3 independent experiments. C) The MFI of mCherry-positive cells from the flow cytometry in (A). Error bars indicate mean ± SD; n = 3 independent experiments. D) qPCR analysis of total viral DNA (HIV) and unintegrated DNA (2-LTR circles) from cells collected in (A). Error bars indicate mean ± SD; n = 3 independent experiments. E) iK562-Scr and iK562-H1-KD cells were treated with Dox and infected with pNCA-mCherry viruses carrying either IN-WT or IN-D184A, or treated with medium alone as a mock control. Cells were analyzed by flow cytometry at 48 h post-infection, and representative histograms of mCherry fluorescence are shown. F) Quantification of mCherry-positive cells from the flow cytometry in (D). Numbers above the bars indicate the fold difference in iK562-Scr relative to iK562-H1-KD. Error bars indicate mean ± SD; n = 3 independent experiments. G) The MFI of mCherry-positive cells from the flow cytometry in (E). Error bars indicate mean ± SD; n = 3 independent experiments. H) qPCR analysis of total viral DNA and 2-LTR circles from cells collected in (D). Error bars indicate mean ± SD; n = 3 independent experiments.

## Discussion

In this study, we demonstrate that the expression of integration-deficient retroviruses, including HIV-1 and MLV, is strongly suppressed in wild-type K562 cells consistent with previous studies (14–16). However, knockdown of H1 specifically alleviated the silencing of HIV-1 unintegrated DNAs, restoring viral expression to levels comparable to those of wild-type integration-competent virus. Notably, unspliced HIV-1 reporter transcripts were strongly upregulated in H1-depleted cells, while spliced reporter transcripts were expressed at approximately half the wild-type level. In contrast, MLV expression, derived from both spliced and unspliced reporter transcripts, remained equally repressed in H1 knockdown and control cells, indicating that H1 depletion selectively impacts HIV-1 but not MLV silencing, and that the restoration of viral expression is not dependent on splicing.

It is well established that linker histone H1 plays important roles in regulating chromatin structure and gene expression. Increasing evidence indicates that H1 is involved in regulating core histone post-translational modifications. For example, a previous study showed that H1 silences repetitive DNA sequences in mouse embryonic stem cells by promoting repressive histone H3K9 methylation and chromatin compaction (36). H1 also enhances deposition of repressive H3K27 methylation by stimulating PRC2 activity and preferentially binding H3K27me3 nucleosomes, establishing a reinforcing feedback loop (34, 37, 38). Conversely, H1 has been shown to block the deposition of H3 post-translational modifications (H3K4 and H3K36 methylations) associated with active gene transcription, both *in vivo* and *in vitro* (33, 34). Mechanistically H1 may restrict H3K4 methylation by blocking access to H3 tails and limiting H3 tail mobility (33, 39). H1 also represses histone acetylation by blocking histone acetyltransferase (HAT) access and by interacting with histone deacetylases (HDACs) (40, 41). This is consistent with observations that treatment with the HDAC inhibitor trichostatin A (TSA) restored expression of unintegrated MLV DNAs, with little effect on wild-type integrated virus, indicating that histone acetylation is a key determinant of unintegrated DNA silencing (Fig 1A and (14).) Collectively, these findings underscore the broad contribution of H1 to chromatin compaction and transcriptional silencing.

Our results suggest that H1 may acts on HIV unintegrated DNAs through mechanisms similar to those observed at host chromatin, through influencing DNA accessibility or modulation of histone epigenetic modifications. By contrast, MLV repression appears to depend more on H1-independent host pathways. For example, the histone chaperones CHAF1A and CHAF1B are essential for silencing unintegrated HIV-1 DNAs by promoting H3K9 trimethylation, but they have no effect on MLV unintegrated DNA, indicating that MLV relies on alternative factors to modify histone methylation (19). This distinction underscores that H1 plays a central role in silencing HIV-1 unintegrated DNA, whereas MLV repression is maintained by different host mechanisms. The basis for the distinction between the responses of the viral genomes to H1 depletion is not clear.

Interestingly, depletion of H1.2, H1.3, H1.4, and H1.5 triggered a strong compensatory increase in H1.x expression. Although less well characterized, H1.x exhibits a H1 binding profile with depletion at active transcription start sites, but unlike other variants, it is enriched at RNAP II– positive sites within gene bodies, suggesting potential roles in supporting ongoing transcription (42). Compensation among H1 subtypes has been reported in multiple systems; for example, H1 triple-knockout (H1c/H1d/H1e) mouse embryonic stem cells display compensatory upregulation of H1.0 and H1.x (43). Whether the compensatory increase of H1.x contributes to maintaining MLV silencing while permitting HIV-1 derepression remains an open question that warrants further investigation

Overall, our findings reveal a novel and selective role of H1 in the silencing of unintegrated HIV-1 DNA. Future studies should dissect the molecular basis for the differential requirement of H1 in HIV-1 versus MLV silencing and explore whether H1 variant compensation or cooperation with specific host factors underlies these distinct outcomes.

## Materials and Methods

### Cells and Plasmids

Human HeLa cells and human Lenti-X 293 T cells (Takara) were cultured in Dulbecco’s Modified Eagle Media (DMEM), supplemented with 10% heat-inactivated fetal bovine serum (FBS), 100 U/ml penicillin, and 100 μg/mL streptomycin. iK562 cells were cultured in Iscove’s Modified Dulbecco’s Medium (IMDM), supplemented with 10% heat-inactivated fetal bovine serum, 100 U/ml penicillin, 100 μg/ml streptomycin and 2 mM L-glutamine.

To generate HIV-1 reporter particles, we used the plasmid pNL4-3.mCherry.R^−^.E^−^ (wt) and its integration-deficient mutant pNL4-3.mCherry.R^−^.E^−^ (IN-D64A), which carries the D64A substitution in the integrase active site. These constructs express HIV-1 Gag-Pol from the plasmid backbone and produce HIV-1 particles expressing mCherry from spliced mRNA (Fig. S1A). We also employed pLVX-mCherry (Fig. S1B), together with the helper constructs Δ8.2 (wt) or Δ8.2 (D84A), which encode either wild-type or integrase-deficient HIV-1 Gag-Pol, to generate HIV-1 particles expressing mCherry from unspliced mRNA. For the Moloney MLV system, we used two reporter plasmids: MLV-mCherry (spliced, Fig. S1C) and pNCA-mCherry (unspliced, Fig. S1D). In addition, constructs expressing Gag-Pol from NB-tropic MLV pCMV-intron (wt) and pCMV-intron (D184A) were used to generate either integration-competent or integration-deficient MLV particles. All reporter viruses were designed as replication-defective, single-round vectors and were pseudotyped with the vesicular stomatitis virus G (VSV-G) envelope, encoded by pMD2.G (a gift from Didier Trono, École Polytechnique Fédérale de Lausanne; Addgene plasmid #12259).

### Histone H1 knock-down

Transient H1 knockdown (H1-KD) cells were generated using a CRISPR interference (CRISPRi) system. A K562 cell line stably expressing high levels of a catalytically inactive Cas9 protein fused to the KRAB repressor domain and mCherry (dCas9-KRAB-mCherry) was transfected with vectors expressing sgRNAs targeting H1.5 (GGCAGGAGCGGTTTCCGACA), H1.2 (GGCTGCCGCCGGCTATGATG), H1.3 (GGCTGCCGCCGGCTATGATG), and H1.4 (GGCCAAGCCTAAGGCTAAAA) (collectively referred to as 4×H1-sgRNA), or with a non-targeting control sgRNA (“scramble” or scr-sgRNA). Following electroporation, cells were selected with puromycin for 96 hours and then harvested. The guide RNAs for each H1 subtype were previously optimized empirically (44). To establish a doxycycline (Dox)-induced H1 knockdown system, K562 cells expressing a dox-inducible dCas9-KRAB-P2A-mCherry were generated by lentiviral transduction with the TET-ON vector pAAVS1-NDi-CRISPRi (addgene #73497). Transduced cells were selected with 200 ug/ml G418 and inducible dCas9-KRAB-P2A-mCherry were further selected with fluorescence-activated cell sorting (FACS) after 3 days of Doxycycline treatment (1ug/ml). Selected cells were allowed to grow in the absence of doxycycline and then transduced with pU6-sgRNA EF1Alpha-puro-T2A-BFP (addgene #60955) which was engineered with 4 tandem U6-sgRNA expression cassettes, each expressing 4xH1-sgRNA. As control, cells were also transduced with a non-targeting pU6-sgRNA EF1Alpha-puro-T2A-BFP (GCACTACCAGAGCTAACTCA). Cells expressing constitutive high levels of sgRNAs were selected by combining puromycin selection (10ug/ml) and FACS to collect the top BFP positive cells. H1 depletion was induced by supplementing culture media with doxycycline 1ug/ml for 5 days and H1 content assayed by RP-HPLC of acid extracted histones.

### Retroviral Particle Production and Infection

We generated eight types of retroviral reporter particles, including both HIV-1 and MLV, spliced and unspliced reporters, and integration-competent or integration-deficient variants. To produce virus, Lenti-X 293T cells (5 × 10^6^ per 10-cm dish) were seeded one day before transfection. Transfections were performed using Lipofectamine 3000 (Thermo Fisher Scientific) according to the manufacturer’s instructions. For the HIV-1 spliced reporter, cells were transfected with 15 μg of pNL4-3.mCherry.R−.E− containing either wild-type integrase (IN-WT) or the integrase mutant (IN-D64A), together with 5 μg of pMD2.G encoding the VSV-G envelope. For the HIV-1 unspliced reporter, cells received 10 μg of pLVX-mCherry, 10 μg of Δ8.2 (IN-WT or IN-D64A), and 4 μg of pMD2.G. For MLV particles, cells were transfected with 8 μg of either MLV-mCherry (spliced) or pNCA-mCherry (unspliced), along with 4 μg of pCMV-intron (IN-WT or IN-D184A) and 4 μg of pMD2.G. Viral supernatants were collected 48 h after transfection, clarified by centrifugation, filtered through a 0.45 μm membrane, treated with TURBO DNase (Invitrogen) for 1 h at 37 °C to remove contaminating plasmid DNA, aliquoted, and stored at −80 °C.

For infection experiments, scrambled control iK562 (iK562-Scr) and histone H1 knockdown iK562 (iK562-H1-KD) cells were seeded into T25 flasks and treated with 3 μg/mL doxycycline (Dox) for 5 days to induce knockdown prior to infection. Equal volumes of viral supernatant from IN-WT and integrase-deficient (IN-D64A or IN-D184A) preparations, produced in parallel, were applied to cells for side-by-side comparison. Infections were carried out at a multiplicity of infection (MOI) ≤ 0.5, which reproducibly yielded 30–50% mCherry-positive cells after 48 h, as measured by flow cytometry.

### Western Blotting

Cells were harvested at the indicated time points after infection, and histones were extracted using the Histone Extraction Kit (Abcam) according to the manufacturer’s instructions. Purified histones were mixed with 4× Laemmli protein loading buffer, boiled at 95 °C for 5 min, separated on 4– 20% Mini-PROTEAN® TGX™ Precast Protein Gels (Bio-Rad), and transferred to nitrocellulose membranes (Bio-Rad). Membranes were blocked with 5% non-fat milk in PBS containing 0.1% Tween-20 (PBST) for 1 h at room temperature (RT) and incubated overnight at 4 °C with primary antibodies against Histone H1.2 (#ab4086, Abcam, 1:1,000), Histone H1.3 (#ab183736, Abcam, 1:1,000), Histone H1.4 (22H6L6, #702876, Thermo Fisher Scientific, 1:1,000), Histone H1.5 (16HCLC, #711912, Abcam, 1:1,000), or Histone H1.x (#ab31972, Abcam, 1:1,000). Protein detection was performed using SuperSignal™ West Femto Maximum Sensitivity Substrate (Thermo Fisher Scientific) and imaged with the ChemiDoc MP Imaging System (Bio-Rad). After signal acquisition, membranes were stripped with ReBlot Plus Strong Antibody Stripping Solution (Millipore) for 30 min at RT, re-blocked in 5% milk in PBST for 1 h, and subsequently incubated overnight at 4 °C with anti-Histone H3 (#ab1791, Abcam, 1:5,000) as an internal loading control. The following day, membranes were incubated for 1 h at RT with HRP-conjugated secondary antibodies (GE Healthcare; NA934V, NA9310V, 1:10,000 in 5% milk/PBST), and signals were developed and imaged as described above.

### RP-HPLC

Histone proteins were isolated by extraction with 0.2N sulfuric acid, as previously described (34, 45, 46). Acid extracted histones were analyzed by electrophoresis through a 15% SDS-polyacrylamide gel and visualized by Coomassie brilliant blue staining. For quantification, acid extracted histones were analyzed by reversed-phase high pressure liquid chromatography with a Waters 2695 system equipped with a Vydac 218TP C18 HPLC column. The effluent was monitored and peaks recorded using the Waters 996 Photodiode Array Detector at 214nm. H1 peak integrations were performed using the Waters Empower Pro software (v.2) and normalized to H2B peaks.

### Flow Cytometry

At 48 h post-infection, iK562 cells were harvested and resuspended in phosphate-buffered saline (PBS) supplemented with 1% heat-inactivated fetal bovine serum (FBS). Flow cytometry was performed on a BD LSRFortessa cytometer (BD Biosciences), and data were analyzed using FlowJo software. Cell populations were first gated for viability based on side scatter area (SSC-A) versus forward scatter area (FSC-A). Single cells were then selected from this viable population using forward scatter height (FSC-H) versus forward scatter area (FSC-A). Finally, mCherry-positive cells were identified and quantified by gating the single-cell population using forward scatter area (FSC-A) versus PE-Texas Red-A fluorescence.

### Viral DNA Detection

Cells were harvested 48 h after infection, and total DNA was extracted using the QIAwave DNA Blood & Tissue Kit (Qiagen) following the manufacturer’s instructions. To quantify viral replication intermediates, 100 ng of DNA was used as template in reactions containing SYBR Green PCR Master Mix (Roche) and 12 pmol of the indicated primers (Table S1). Reactions were performed in 96-well plates on a 7900 Fast Real-Time PCR System (Applied Biosystems) under the following cycling conditions: 50 °C for 2 min, 95 °C for 10 min, followed by 40 cycles of 95 °C for 15 s, 60 °C for 30 s, and 72 °C for 30 s. A melting curve analysis was carried out at the end of each run to verify amplification specificity, according to the manufacturer’s recommendations. mCherry primers were used to detect total viral DNA, 2-LTR primers were used to detect circular viral DNA containing two tandem long terminal repeats (LTRs), and LTR primers were used to detect all forms of circular viral DNA. Threshold cycle (Ct) values were normalized to GAPDH, and relative DNA levels were calculated using the 2^ΔCt^ method.

## Statistical Analysis

Data are presented as mean ± standard deviation (SD). Statistical methods and sample sizes are described in the corresponding figure legends. Comparisons between two groups were performed using an unpaired Student’s t-test, with Welch’s correction applied when variances were unequal. Significance levels were defined as: ns (P ≥ 0.05), * (P < 0.05), ** (P < 0.01), and *** (P < 0.001).

## Acknowledgement

H.D.P., S.E.H., L.N.M. and A.I.S. were supported by NIH grant R01 GM147165. J.Z., G.Z.W. and S.P.G. were supported by NIH grants R01 CA030488 and U54 AI170660-03.

## Supporting information

Fig. S1: Plasmids maps of reporter viruses

Table S1: Primers for qRT-PCR

## References

1. P. O. Brown, B. Bowerman, H. E. Varmus, J. M. Bishop, Retroviral integration: structure of the initial covalent product and its precursor, and a role for the viral IN protein. Proc Natl Acad Sci U S A 86, 2525–2529 (1989).

2. C. M. Farnet, W. A. Haseltine, Integration of human immunodeficiency virus type 1 DNA in vitro. Proc Natl Acad Sci U S A 87, 4164–4168 (1990).

3. M. D. Miller, C. M. Farnet, F. D. Bushman, Human immunodeficiency virus type 1 preintegration complexes: studies of organization and composition. J Virol 71, 5382–5390 (1997).

4. T. Roe, T. C. Reynolds, G. Yu, P. O. Brown, Integration of murine leukemia virus DNA depends on mitosis. EMBO J 12, 2099–2108 (1993).

5. P. F. Lewis, M. Emerman, Passage through mitosis is required for oncoretroviruses but not for the human immunodeficiency virus. J Virol 68, 510–516 (1994).

6. J. B. Weinberg, T. J. Matthews, B. R. Cullen, M. H. Malim, Productive human immunodeficiency virus type 1 (HIV-1) infection of nonproliferating human monocytes. J Exp Med 174, 1477–1482 (1991).

7. P. Lewis, M. Hensel, M. Emerman, Human immunodeficiency virus infection of cells arrested in the cell cycle. EMBO J 11, 3053–3058 (1992).

8. M. Yamashita, M. Emerman, Capsid is a dominant determinant of retrovirus infectivity in nondividing cells. J Virol 78, 5670–5678 (2004).

9. K. A. Matreyek, A. Engelman, Viral and cellular requirements for the nuclear entry of retroviral preintegration nucleoprotein complexes. Viruses 5, 2483–2511 (2013).

10. F. K. Yoshimura, R. A. Weinberg, Restriction endonuclease cleavage of linear and closed circular murine leukemia viral DNAs: discovery of a smaller circular form. Cell 16, 323–332 (1979).

11. J. M. Kilzer et al., Roles of host cell factors in circularization of retroviral dna. Virology 314, 460–467 (2003).

12. P. Schwartzberg, J. Colicelli, S. P. Gob, Construction and analysis of deletion mutations in the pol gene of Moloney murine leukemia virus: a new viral function required for productive infection. Cell 37, 1043–1052 (1984).

13. H. Sakai et al., Integration is essential for ebicient gene expression of human immunodeficiency virus type 1. J Virol 67, 1169–1174 (1993).

14. G. Z. Wang, Y. Wang, S. P. Gob, Histones Are Rapidly Loaded onto Unintegrated Retroviral DNAs Soon after Nuclear Entry. Cell Host Microbe 20, 798–809 (2016).

15. F. K. Geis, S. P. Gob, Unintegrated HIV-1 DNAs are loaded with core and linker histones and transcriptionally silenced. Proc Natl Acad Sci U S A 116, 23735–23742 (2019).

16. F. K. Geis, D. P. Kelenis, S. P. Gob, Two lymphoid cell lines potently silence unintegrated HIV-1 DNAs. Retrovirology 19, 16 (2022).

17. W. M. Schneider, D. T. Wu, V. Amin, S. Aiyer, M. J. Roth, MuLV IN mutants responsive to HDAC inhibitors enhance transcription from unintegrated retroviral DNA. Virology 426, 188–196 (2012).

18. Y. Zhu, G. Z. Wang, O. Cingoz, S. P. Gob, NP220 mediates silencing of unintegrated retroviral DNA. Nature 564, 278–282 (2018).

19. F. K. Geis et al., CHAF1A/B mediate silencing of unintegrated HIV-1 DNAs early in infection. Proc Natl Acad Sci U S A 119 (2022).

20. L. Dupont et al., The SMC5/6 complex compacts and silences unintegrated HIV-1 DNA and is antagonized by Vpr. Cell Host Microbe 29, 792–805 e796 (2021).

21. I. D. Irwan, H. P. Bogerd, B. R. Cullen, Epigenetic silencing by the SMC5/6 complex mediates HIV-1 latency. Nat Microbiol 7, 2101–2113 (2022).

22. K. Luger, A. W. Mader, R. K. Richmond, D. F. Sargent, T. J. Richmond, Crystal structure of the nucleosome core particle at 2.8 A resolution. Nature 389, 251–260 (1997).

23. R. D. Kornberg, Chromatin structure: a repeating unit of histones and DNA. Science 184, 868–871 (1974).

24. S. P. Hergeth, R. Schneider, The H1 linker histones: multifunctional proteins beyond the nucleosomal core particle. EMBO Rep 16, 1439–1453 (2015).

25. W. F. Marzlub, Metazoan replication-dependent histone mRNAs: a distinct set of RNA polymerase II transcripts. Curr Opin Cell Biol 17, 274–280 (2005).

26. D. V. Fyodorov, B. R. Zhou, A. I. Skoultchi, Y. Bai, Emerging roles of linker histones in regulating chromatin structure and function. Nat Rev Mol Cell Biol 19, 192–206 (2018).

27. J. Zlatanova, D. Doenecke, Histone H1 zero: a major player in cell diberentiation? FASEB J 8, 1260–1268 (1994).

28. N. Happel, E. Schulze, D. Doenecke, Characterisation of human histone H1x. Biol Chem 386, 541–551 (2005).

29. B. Drabent, E. Kardalinou, D. Doenecke, Structure and expression of the human gene encoding testicular H1 histone (H1t). Gene 103, 263–268 (1991).

30. I. Martianov et al., Polar nuclear localization of H1T2, a histone H1 variant, required for spermatid elongation and DNA condensation during spermiogenesis. Proc Natl Acad Sci U S A 102, 2808–2813 (2005).

31. M. Tanaka, J. D. Hennebold, J. Macfarlane, E. Y. Adashi, A mammalian oocyte-specific linker histone gene H1oo: homology with the genes for the oocyte-specific cleavage stage histone (cs-H1) of sea urchin and the B4/H1M histone of the frog. Development 128, 655–664 (2001).

32. C. L. Woodcock, A. I. Skoultchi, Y. Fan, Role of linker histone in chromatin structure and function: H1 stoichiometry and nucleosome repeat length. Chromosome Res 14, 17–25 (2006).

33. S. M. Yang, B. J. Kim, L. Norwood Toro, A. I. Skoultchi, H1 linker histone promotes epigenetic silencing by regulating both DNA methylation and histone H3 methylation. Proc Natl Acad Sci U S A 110, 1708–1713 (2013).

34. M. A. Willcockson et al., H1 histones control the epigenetic landscape by local chromatin compaction. Nature 589, 293–298 (2021).

35. W. C. Goh et al., HIV-1 Vpr increases viral expression by manipulation of the cell cycle: a mechanism for selection of Vpr in vivo. Nat Med 4, 65–71 (1998).

36. S. E. Healton et al., H1 linker histones silence repetitive elements by promoting both histone H3K9 methylation and chromatin compaction. Proc Natl Acad Sci U S A 117, 14251–14258 (2020).

37. C. Martin, R. Cao, Y. Zhang, Substrate preferences of the EZH2 histone methyltransferase complex. J Biol Chem 281, 8365–8370 (2006).

38. J. M. Kim et al., Linker histone H1.2 establishes chromatin compaction and gene silencing through recognition of H3K27me3. Sci Rep 5, 16714 (2015).

39. A. Stutzer et al., Modulations of DNA Contacts by Linker Histones and Post-translational Modifications Determine the Mobility and Modifiability of Nucleosomal H3 Tails. Mol Cell 61, 247–259 (2016).

40. J. E. Herrera, K. L. West, R. L. Schiltz, Y. Nakatani, M. Bustin, Histone H1 is a specific repressor of core histone acetylation in chromatin. Mol Cell Biol 20, 523–529 (2000).

41. A. Vaquero et al., Human SirT1 interacts with histone H1 and promotes formation of facultative heterochromatin. Mol Cell 16, 93–105 (2004).

42. R. Mayor et al., Genome distribution of replication-independent histone H1 variants shows H1.0 associated with nucleolar domains and H1X associated with RNA polymerase II-enriched regions. J Biol Chem 290, 7474–7491 (2015).

43. Y. Fan et al., Histone H1 depletion in mammals alters global chromatin structure but causes specific changes in gene regulation. Cell 123, 1199–1212 (2005).

44. L. A. Gilbert et al., Genome-Scale CRISPR-Mediated Control of Gene Repression and Activation. Cell 159, 647–661 (2014).

45. Q. Lin et al., Reductions in linker histone levels are tolerated in developing spermatocytes but cause changes in specific gene expression. J Biol Chem 279, 23525–23535 (2004).

46. A. M. Sirotkin et al., Mice develop normally without the H1(0) linker histone. Proc Natl Acad Sci U S A 92, 6434–6438 (1995).

