## Supplementary Fig S1 and Table S1 for "Histone H1 Promotes Silencing of Unintegrated HIV-1 DNA"

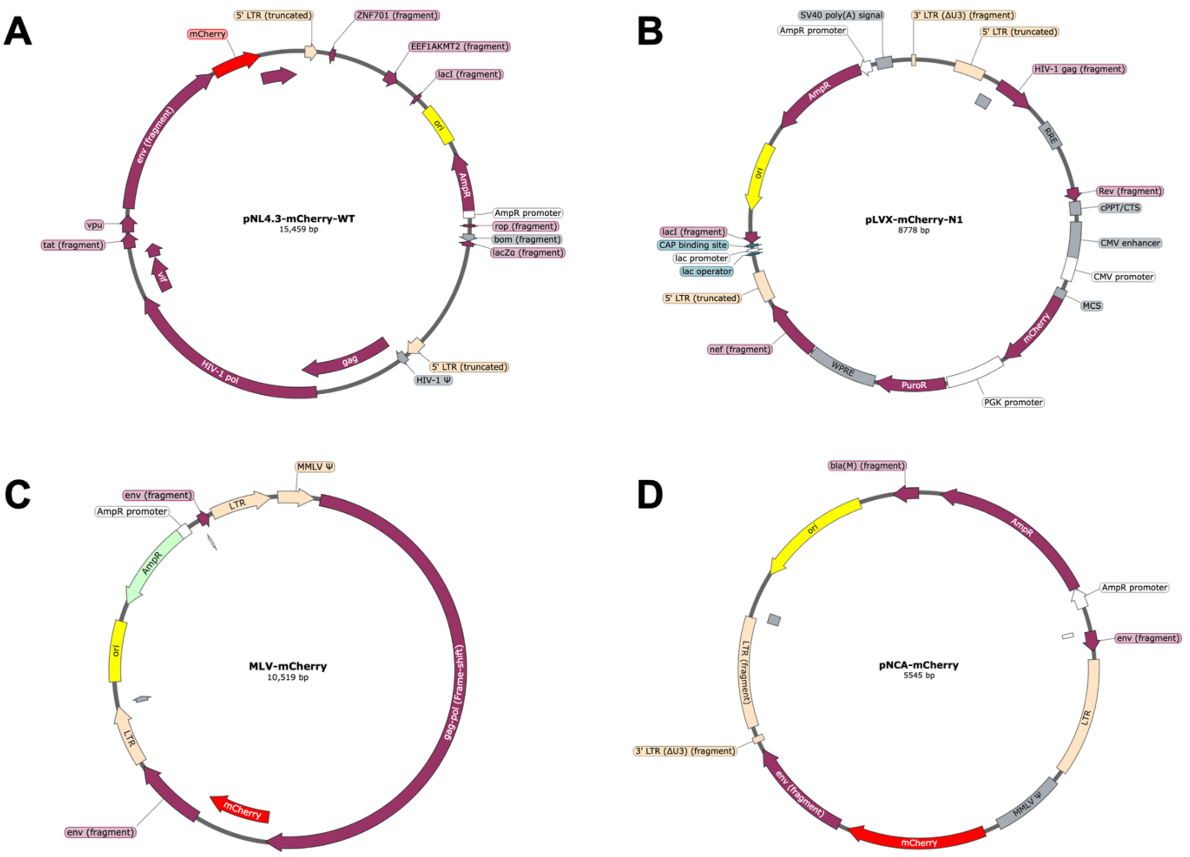


**Fig S1.** Plasmids maps of reporter viruses

A) Plasmids map of pNL4-3.mCherry.R⁻.E⁻

B) Plasmids map of pLVX-mCherry

C) Plasmids map of MLV-mCherry

D) Plasmids map of pNCA-mCherry

**Table S1. Primers for qRT-PCR**

| Target | | Sequence (5' → 3') |
|---|---|---|
| mCherry | Forward | CACGAGTTCGAGATCGAGGG |
| | Reverse | CAAGTAGTCGGGGATGTCGG |
| HIV 2-LTR | Forward | AACTAGGGAACCCACTGCTTAAG |
| | Reverse | TCCACAGATCAAGGATATCTTGTC |
| MLV 2-LTR | Forward | AGGGTCTCCTCTGAGTGATT |
| | Reverse | ATGGTGTGTGGAGGAGTATAAAG |
| MLV LTR | Forward | AGTCCTCCGATTGACTGAG |
| | Reverse | CTCTTTTATTGAGCTCGGG |
| GAPDH | Forward | CAATTCCCCATCTCAGTCGT |
| | Reverse | TAGTAGCCGGGCCCTACTTT |
